# Diminished and altered cellular senescence response in delayed wound healing of aging

**DOI:** 10.1101/2025.06.08.658533

**Authors:** Maria Shvedova, Rex Jeya Rajkumar Samdavid Thanapaul, Qiaoling Wang, Grace Haeun Shin, Jannat Dhillon, Daniel Sam Roh

## Abstract

The transient upregulation of cellular senescence within wound tissues has been demonstrated to be an important biological process facilitating efficient tissue repair. Dysregulation of this transient wound-induced senescence-like response can result in impaired healing outcomes. Given the established age-related decline in tissue regenerative capacity, we hypothesized that alterations in this senescence response contribute to the delayed healing of cutaneous wounds in aged individuals. Our investigation demonstrated a significant delay in the closure of full-thickness dorsal skin wounds in aged mice compared to their young counterparts. Analysis of the wound microenvironment revealed a transient upregulation of senescence-associated markers (p16, p21, senescence-associated β-galactosidase) and senescence-associated secretory phenotype factors in the wound tissue of young mice, a response that was markedly attenuated in aged mice. Single-cell RNA sequencing analysis of all cells isolated from day 6 wounds identified a distinct population of p16^+^/p21^+^/Ki67^-^ senescent fibroblasts in young mice, characterized by a transcriptional signature indicative of pro-healing extracellular matrix production, a finding corroborated in human wound tissue from young donors. Crucially, in aged wounds, we observed a lower quantity of these senescent cells, a deficit compounded by a qualitative, age-dependent shift in their function, moving away from beneficial extracellular matrix remodeling towards a more detrimental pro-inflammatory state, which ultimately can contribute to the delayed wound healing.

## Introduction

Impaired wound healing and persistent and chronic wounds in older individuals represent increasing clinical and economic challenge worldwide^1^. Aging contributes to heightened susceptibility to injury and delayed wound healing, influenced by changes in cellular functions and composition^2^. Senescent cells can adversely affect both the local tissue microenvironment and the entire organism^3^. However, transient upregulation of acute cellular senescence after cutaneous wounding plays an important physiological role during tissue repair and regeneration^4–6^. A significant mechanism through which senescent cells impact surrounding cells is the production of the senescence-associated secretory phenotype (SASP), comprised of inflammatory mediators, extracellular matrix (ECM) modifiers, and growth factors^7^. An example of beneficial upregulation of senescence signaling was demonstrated in zebrafish, where transient senescent cells at the fin amputation site supported tissue regeneration, and their selective elimination impaired healing^4^. In young 2-month-old mice, transiently upregulated senescence signaling during cutaneous wound healing proved to be an important physiological mechanism, facilitating the wound healing process^8^. Elimination of these senescent cells delayed wound healing in young mice^8^.

Chronic accumulation of senescent cells is a well-established hallmark of aging and contributes to tissue dysfunction and impaired regeneration^6^. In contrast, acute and transient senescence upregulation, a short-lived and tightly regulated process arising in response to injury, plays a beneficial role in tissue repair, particularly in young organisms which appears evolutionarily conserved across multiple species^4,8–11^. However, it remains unclear whether this reparative senescence program is preserved with aging across species. While aged tissues exhibit an increased burden of chronic senescent cells^12,13^, whether aging impairs the induction, resolution, or functional identity of acute wound-associated senescent cells is not well understood. Elucidating how aging alters the dynamics and roles of acute senescent cells in the wound environment is important for understanding the mechanisms underlying delayed healing in older adults and for guiding therapeutic development.

We aimed to define the acute senescence response at single-cell resolution in young versus aged mice, focusing on senescence marker expression, SASP profiles, and senescent cell type composition. By characterizing the functional features and prevalence of senescent cell populations during wound healing, we sought to uncover age-associated alterations in this transient program that may contribute to the impaired tissue repair in older adults.

## Materials and methods

### Animals and surgical procedure

Young (2-month-old) and old (24-month-old) C57BL/6 wild type male mice were obtained from Jackson laboratory and National Institute on Aging (NIA), respectively. We utilized single 1-cm full-thickness excisional cutaneous wound model on the dorsal midline under isoflurane anesthesia. Wounds were covered with Tegaderm^TM^ (3M), which was changed every 3 days. Mice were given extended-release buprenorphine subcutaneously (0.03 mg/kg) for analgesia and housed with food and water ad libitum. Mice were monitored postoperatively every three days with digital imaging. Wound area was calculated using ImageJ software. Mice from each group were euthanized at days 6, 12, 18, and 24 (at least 5 animals per group per time point), and the wound tissue was harvested. Skin obtained during the initial wound creation was used as a non-wounded control. All procedures were performed in accordance with the recommendations outlined in the Guide for the Care and Use of Laboratory Animals of the National Institutes of Health and were approved by the Boston University Subcommittee on Research and Animal Care.

### Senescence-associated β-galactosidase analysis

Analysis of senescence-associated β-galactosidase (SA-β-gal) during wound healing was performed using CellEvent^tm^ Senescence Green Detection Kit (Invitrogen, cat. #C10851). After fixation in 4% paraformaldehyde in PBS for 2 h at 4 °C with subsequent overnight sucrose treatment at 4 °C, the 5-μm-thick cryosections were washed in PBS, traced with hydrophobic pen, followed by washing in 1% BSA in PBS; then, 100 μl of prewarmed at 37 °C working solution was added to each section. Sections were incubated overnight at 37 °C without CO2 in humidified chamber in the dark. After incubation, working solution was removed, and sections were washed three times with PBS, counterstained with DAPI, mounted, and coverslipped. Fluorescent microscopy images were captured using Nikon Eclipse E400 fluorescent microscope at 10x magnification. Percentage of SA-β-gal-positive cells in the dermis and epidermis was calculated using ImageJ software. Hair follicles, sebaceous glands, and adipose skin layer were excluded from SA-β-gal analysis as they consistently stained positive.

### Immunostaining

For immunostaining, after fixation in 4% paraformaldehyde in PBS for 2 h at 4 °C with subsequent overnight sucrose treatment at 4 °C, the 5-μm-thick cryosections were washed in PBS. Antigen retrieval was performed in EnVision FLEX Target Retrieval Solution Low pH (Dako, ref. #K8005) using Dako PT Link antigen retrieval machine. Sections were washed with PBS and traced with hydrophobic pen. Blocking was performed for 1 h at room temperature with 10% horse serum in PBS with 0.05% Triton X-100. Sections were incubated with rabbit anti-p21 primary antibodies (ab188224) overnight at 4 °C (1:100), washed with PBS, followed by incubation with secondary donkey antirabbit antibodies conjugated with Alexa Fluor® 594 (ab150064) for 1h at RT (1:1000). After aspiration of secondary antibodies and extensive washing to remove unbound antibodies, sections were counterstained with DAPI, mounted, and coverslipped. Fluorescent microscopy images were captured using Nikon Eclipse E400 fluorescent microscope at 20x magnification. Percentage of p21-positive cells in the dermis and epidermis was calculated using ImageJ software.

### Gene expression analysis

Harvested skin tissue was bulk analyzed with qPCR for senescence cell cycle arrest markers p16, p21, and p53, and SASP markers (Il6, Mcp1/Ccl2, Mmp3, Mmp8, Mmp9, Tnf, Tgfb1). Tissue was homogenized using gentleMACS dissociator (Miltenyl Biotec) in TRIzol. RNA extraction was performed using Direct-zol^tm^ RNA MiniPrep Plus kit (Zymo Research, cat. #R2070). After RNA quantification by Nanodrop, cDNA synthesis was performed using Verso cDNA synthesis kit (ThermoScientific, ref. #AB-1453/B). qPCR was performed using Maxima SYBR Green/ROX qPCR Master Mix (ThermoScientific, ref. #K0222) in the StepOnePlus real-time PCR system (Applied Biosystems). Relative quantitative RNA was normalized using the housekeeping gene β-actin. qPCR primers are presented in Table S1.

### Protein analysis

#### Western blotting

Harvested skin was homogenized using gentleMACS dissociator (Miltenyl Biotec) in RIPA Lysis Buffer System with protease inhibitor cocktail (ChemCruz). Protein concentrations were determined using the Coomassie Plus (Bradford) Assay Kit and spectrophotometer. Samples were denatured with Pierce™ Lane Marker Reducing Sample Buffer (Thermo Scientific). Protein electrophoresis was performed loading 20 μg protein from each sample for separation by 10% SDS-PAGE. Protein was transferred to nitrocellulose membranes (Life technologies). Membranes were blocked with Pierce™ Clear Milk Blocking Buffer (Thermo Scientific) and then incubated overnight with the primary antibodies against MMP-9 (ab38898), p21 (ab188224), and GAPDH (ab9485), washed three times for 10 minutes with PBST, followed by incubation with horseradish peroxidase-conjugated secondary antibodies (ab7090). After washing the membrane four times for 10 min with PBST, immobilon Western Chemiluminescent HRP Substrate (Millipore) was added, and the result was tested with ChemiDoc XRS+ imaging system (Bio-Rad Laboratories). Results were analyzed with ImageJ using NIH method (protein expression relative to GAPDH).

#### ELISA

IL-6 expression was confirmed with ELISA using Mouse ELISA Kit (ab100713 – IL-6) in accordance with the manufacturer instructions.

### Statistical analysis

Statistical analysis was performed using two-way ANOVA with factors *age* and *timepoint*, followed by post-hoc tests for multiple comparisons. For comparisons between young and old animals at individual timepoints, two-tailed unpaired t-tests were used. Results are expressed as mean ± SD. p < 0.05 was considered statistically significant.

### Wound digestion

For single-cell sequencing experiment, 1-cm full-thickness dorsal cutaneous wound was created in young (2-month-old) and old (24-month-old) male mice (n=3 per age group, cells were pooled) under isoflurane anesthesia with postoperative care outlined above. Mice were euthanized on day 6 after wounding, and wounds were separated from subcutaneous fat tissue, collected, cut in 2-mm-pieces, digested in shaker for 1h 40min in 0.2% Collagenase solution in DMEM without FBS at 37 C° and 200rpm with vortexing every 20 minutes. After digestion, cell-containing solution was passed through the cell strainer, diluted with complete media (1:1), and spun down. Supernatant was removed, and cells were resuspended in PBS containing 10% FBS, spun down to remove traces of media, resuspended in the same buffer, and used for single cell sequencing.

### Mouse single-cell sequencing data processing

Raw sequencing output (BCL) files were converted to FASTQ using the Cell Ranger mkfastq module (v6.0) and processed with the count pipeline (v6.0) to align reads to the *mm10* mouse reference genome and generate gene-by-barcode UMI matrices. Each sample was processed using Seurat (v5.3.0) in R (v4.4.2). Cells from 2-month-old and 24-month-old mouse samples were retained if they had >200 and <7,500 detected genes (nFeature_RNA) and <10% mitochondrial transcript content. Filtered 2-month and 24-month Seurat objects were normalized using LogNormalize, and highly variable genes were identified. Each dataset was scaled and reduced using PCA before doublet detection with scDblFinder. Singlets were retained and merged into a combined object, which was re-normalized, scaled (regressing out nCount_RNA and percent.mt), and batch-corrected using Harmony on the top 30 principal components. Dimensionality reduction and visualization were performed using Uniform Manifold Approximation and Projection (UMAP) (dims 1–30), and Louvain clustering was applied at resolution 0.5. Cluster markers were identified using presto::wilcoxauc (log₂FC > 0.25, FDR < 0.05) and Seurat::FindAllMarkers (min.pct = 0.1, logfc.threshold = 0.25). Clusters were annotated based on known cell type–specific gene signatures from prior literature.

### Human single-cell sequencing data processing

Publicly available scRNA-seq data from human skin and wound samples (GSE241132) were downloaded from GEO, including raw count matrices and metadata spanning four conditions: Skin (baseline), Wound1 (day 1), Wound7 (day 7), and Wound30 (day 30). Data were processed using Seurat (v5.3.0) in R (v4.4.2) as above. Cells with 200–7,500 detected genes and <10% mitochondrial content were retained. Each sample was normalized, scaled, and subjected to PCA. Doublets were removed using scDblFinder, and singlets were merged into a single Seurat object with metadata added via AddMetaData. To correct for batch effects across conditions, RunHarmony was applied to the top 30 PCs using “condition” as the integration variable. UMAP (dims 1–30) and graph-based clustering (FindNeighbors, FindClusters, resolution = 0.5) were performed on Harmony embeddings to define transcriptionally distinct cell populations. Cluster markers were identified with FindAllMarkers (Wilcoxon test, adj. p < 0.05, log₂FC > 0.25, min.pct = 0.1). Cell types were annotated based on canonical markers

### Gene set enrichment pathway analysis

Genes ranked by log₂ fold-change between senescent (p16⁺/p21⁺/Mki67⁻) and other cells were subjected to Gene Set Enrichment Analysis (GSEA) using the fgseaMultilevel function (fgsea v1.20.0) against MSigDB Hallmark, GO Biological Process, and custom senescence (SAUL_SEN_MAYO) gene sets. Pathways with FDR < 0.05 were deemed significant. Normalized Enrichment Scores (NES) were extracted and plotted as ordered bar charts in ggplot2, colored by enrichment direction.

### CellChat intercellular signaling analysis

Log-normalized RNA counts from the 2-month and 24-month samples were used to construct CellChat objects for intercellular communication analysis. Cells were grouped into three categories: “Senescent_FB” (fibroblast clusters 1, 2, and 15 with *Cdkn2a* > 0, *Cdkn1a* > 0, *Mki67* = 0), “Other_FB” (non-senescent fibroblasts), and “Immune_Cells” (clusters 0, 3, 4, 5, 6, 7, 10, 11, 12). CellChatDB.mouse was used as the reference ligand–receptor database. For each age group, we followed the standard CellChat workflow: subsetData(), identifyOverExpressedGenes(), identifyOverExpressedInteractions(), computeCommunProb(), filterCommunication(min.cells = 10), computeCommunProbPathway(), and aggregateNet(). The resulting CellChat objects were merged using mergeCellChat(), and differential signaling between 2-month and 24-month samples was assessed using netVisual_diffInteraction() (Δ = 2mo – 24mo). Outgoing signals from the “Senescent_FB” group were extracted (subsetCommunication(slot.name = “net”)), summarized by signaling pathway, and visualized using bar and bubble plots to highlight differences in total communication probability between age groups. Aggregate communication probabilities for each cell-type pair were extracted directly from cellchat_2mo@net$weight and cellchat_24mo@net$weight (each a square sender×receiver matrix) after running computeCommunProb, computeCommunProbPathway, and aggregateNet. The difference matrix (Δ = 2 mo − 24 mo) was reshaped with reshape2::melt into a long data frame (Sender, Receiver, DeltaWeight) and plotted as a ggplot2 geom_tile heatmap using a red–white–blue gradient (red = higher in 2 mo, blue = higher in 24 mo).

## Results

### Significant delay in wound healing in aged mice

Evaluation of full-thickness dorsal skin wound closure kinetics revealed a statistically significant delay in aged mice compared to their younger counterparts across the monitored time course (Fig. 1A). The divergence in healing rates became apparent as early as day 6 post-wounding, at which point aged mice exhibited significantly larger open wound surface areas that persisted through subsequent time points (Fig. 1B). Quantification of complete wound closure, a clinically relevant endpoint^14^, demonstrated that while nearly all young mice achieved full epithelialization by day 18, the majority of aged mice required an additional 3 to 6 days for complete wound closure, healing between days 22 and 24 (Fig. 1C). These data establish that aging is associated with a substantial impairment in the rate and completion of cutaneous wound healing in this murine model.

**Figure 1.**
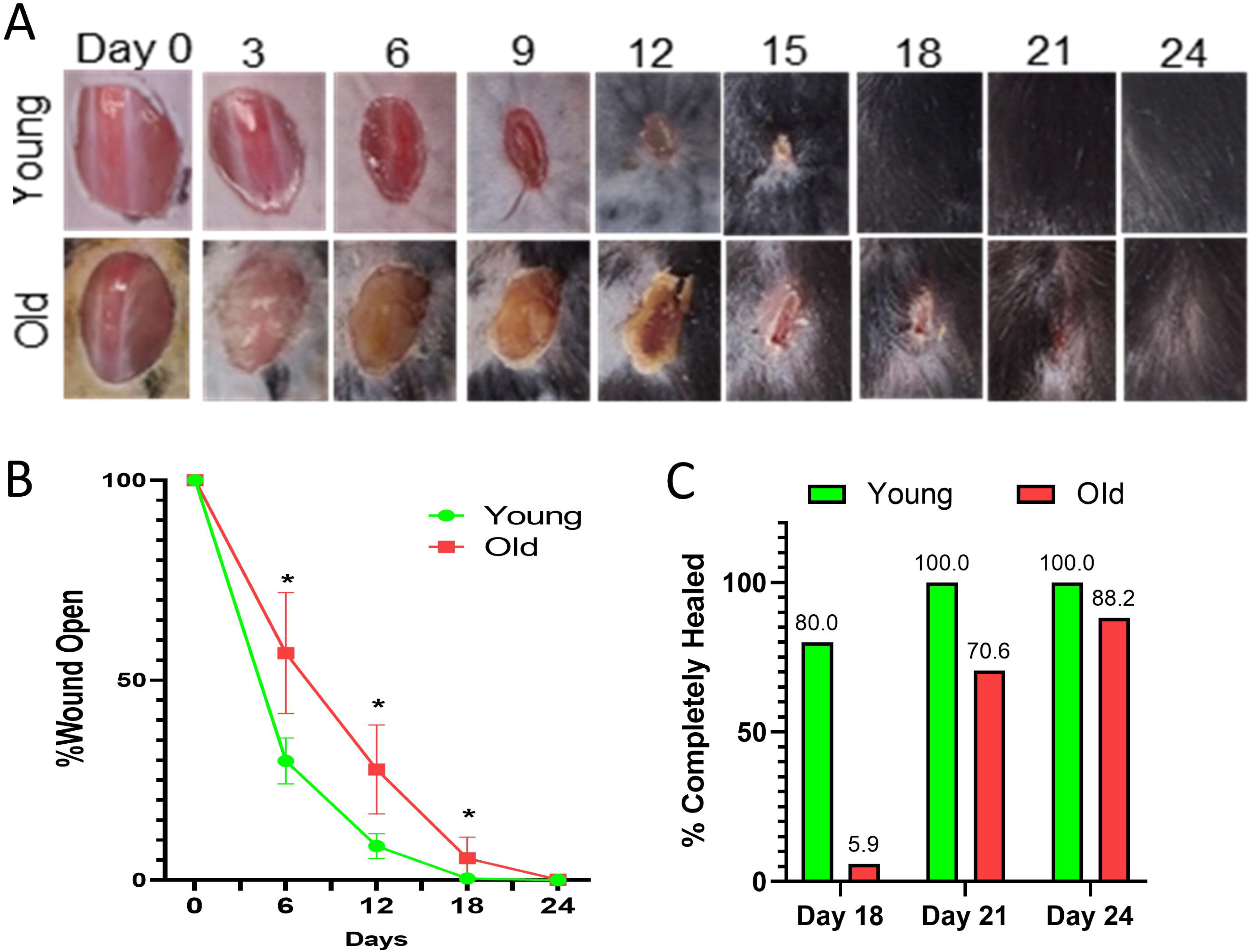
Delayed wound healing in aged mice. (A) Representative macroscopic images of full-thickness dorsal skin wounds in young (2-month-old) and old (23-24-month-old) mice at days 0, 3, 6, 9, 12, 15, 18, 21, and 24 post-wounding. (B) Quantification of the percentage of open wound area over time in young and old mice. N=5 per age group per time point. Asterisks (*) indicate statistically significant differences between young and old mice at the indicated time points (p<0.05). (C) Percentage of young and old mice with completely healed wounds at days 18, 21, and 24 post-wounding.

### Transient upregulation of SA-β-gal activity and p16 expression during wound healing occurs in young but not in old mice

Analysis of SA-β-gal activity, a widely used marker of cellular senescence, demonstrated a striking difference between age groups in wound tissues. Fluorescence staining revealed a robust accumulation of SA-β-gal-positive cells within the granulation tissue of young mice starting from day 6 and diminishing through day 18 (Fig. 2A). Quantification demonstrated a statistically significant increase in SA-β-gal-positive cells in young mice at days 6, 12, and 18 compared to the baseline (Day 0) and compared to the aged mice at the same time points (p<0.05, p<0.01, Student’s t-test) (Fig. 2C). In stark contrast, aged mice exhibited no significant increase in SA-β-gal activity above baseline levels throughout the entire 24-day healing period (Fig. 2B, C).

**Figure 2.**
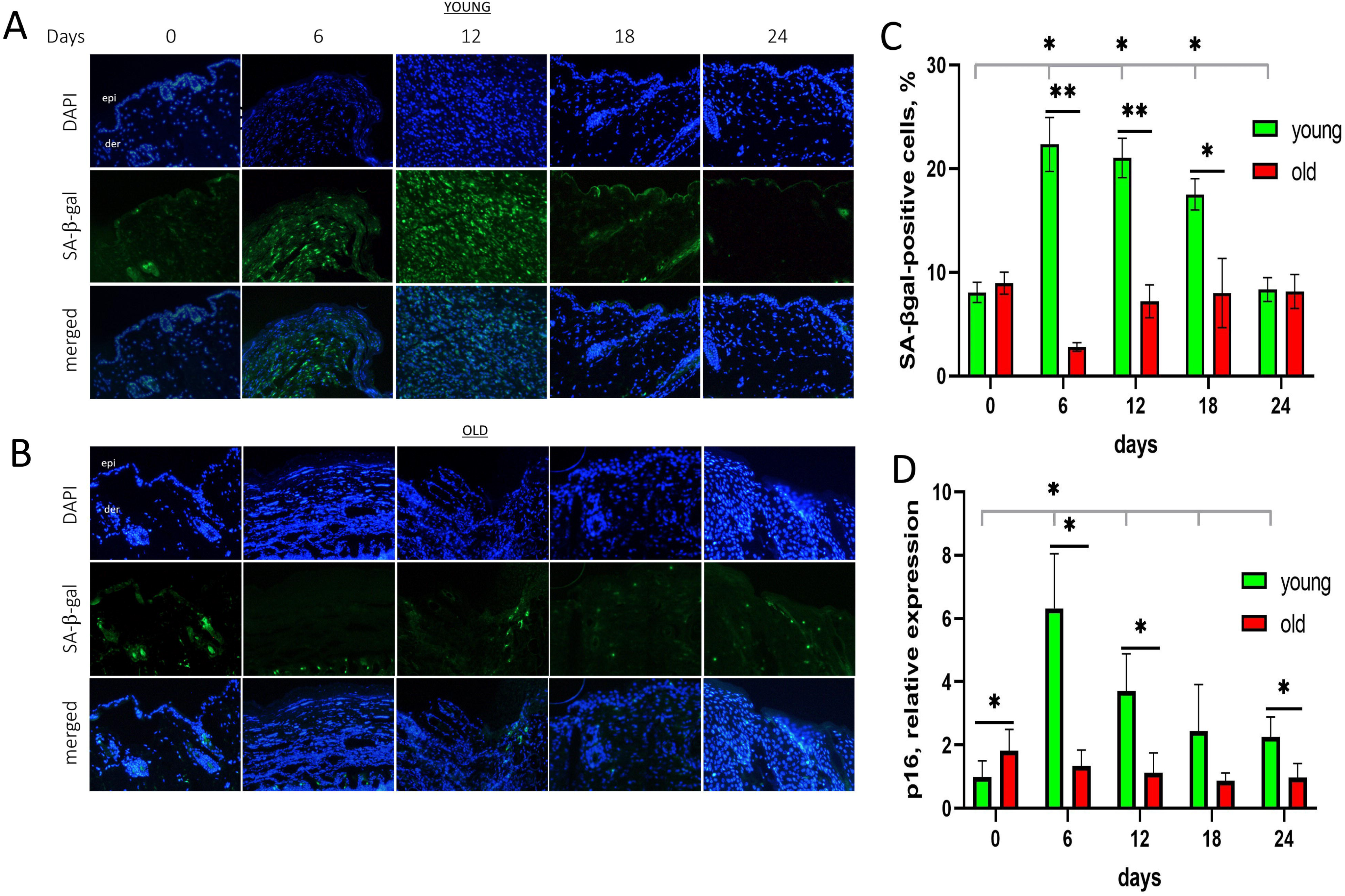
Transient upregulation of SA-β-gal activity and p16 during wound healing in young but not in old mice. (A) Representative immunofluorescence images of wound tissue sections from young mice at days 0, 6, 12, 18, and 24 post-wounding, stained for SA-β-gal activity (green) and counterstained with DAPI (blue) to visualize nuclei. Epi=epidermis, der=dermis for reference. (B) Representative immunofluorescence images of wound tissue sections from old mice at days 0, 6, 12, 18, and 24 post-wounding, stained for SA-β-gal activity (green) and counterstained with DAPI (blue). (C) Quantification of the percentage of SA-β-gal-positive cells in the wound tissue of young and old mice at days 0, 6, 12, 18, and 24 post-wounding. (D) Quantification of p16 mRNA relative expression in the wound tissue of young and old mice at days 0, 6, 12, 18, and 24 post-wounding. N=5 per age group per time point. Asterisks (*) indicate statistically significant differences between young and old mice at the indicated time points (p<0.05). Double asterisks (**) indicate p<0.01.

Similarly, we evaluated the mRNA expression of p16 (Cdkn2a), a critical cell cycle inhibitor frequently upregulated in senescent cells. qRT-PCR revealed significantly elevated baseline p16 mRNA levels in unwounded aged skin compared to young skin (p<0.05) (Fig. 2D). Following wounding, young mice demonstrated a robust and transient increase in p16 mRNA expression, peaking at day 6 (p<0.01 vs Day 0 young; p<0.01 vs aged at Day 6), before returning towards baseline levels by day 18 (Fig. 2D). Consistent with the SA-β-gal data, aged mice did not exhibit any significant wound-induced increase in p16 mRNA expression above their elevated baseline at any time point post-wounding (Fig. 2D).

### Increase in p21 positive cells during the wound healing is evident only in young mice

We next assessed the expression dynamics of p21 (Cdkn1a), another key cyclin-dependent kinase inhibitor associated with senescence and cell cycle arrest. Immunofluorescence staining revealed a transient accumulation of p21-positive cells within the young wound tissue, becoming evident from day 12 and peaking at day 18 post-wounding (Supplementary Fig. 2A). Quantification of p21-positive cells in young mice demonstrated a significant increase at days 12, 18, and 24 compared to baseline and aged mice at the same time points (p<0.05) (Supplementary Fig. 2C). In contrast, aged mice did not exhibit a significant increase in the percentage of p21-positive cells compared to the baseline levels throughout the healing period (Supplementary Fig. 2B) and had lower p21 protein levels in bulk wound lysates (Supplementary Fig. 3A, C). This age-dependent difference in p21 induction was further corroborated by qRT-PCR analysis, which demonstrated a significant increase in p21 mRNA expression in young but not aged wounds at days 18 and 24 (p<0.05, p<0.01) (Supplementary Fig. 2D). Consistent with the p21 dynamics, mRNA expression of its upstream regulator p53 (Trp53) also revealed a transient increase in young mice peaking around day 18, which was absent in aged mice by qRT-PCR (Supplementary Fig. 1A).

### Wound tissue expression of SASP markers significantly differs during young and aged wound healing

Analysis of SASP components in homogenized wound tissue by qRT-PCR, Western blot, and ELISA revealed age-dependent differences in their expression profiles. In young mice, multiple pro-inflammatory cytokines and matrix-remodeling factors associated with the SASP, including Il6, Mcp1/Ccl2, Mmp3, Mmp9, Tnf, and Tgfb1, demonstrated transcriptional upregulation peaking primarily around day 6 and returning to baseline by day 12 (Supplementary Fig. 1B-H). Western blot analysis confirmed transient increase in MMP-9 protein levels (Supplementary Fig. 3A and B) and ELISA confirmed increase in IL-6 (Supplementary Fig. 3D) in homogenized young wound tissue.

The SASP component expression in aged wound tissue was significantly attenuated and dysregulated. While IL-6 mRNA and protein were upregulated in aged wounds at day 6, this elevation was prolonged compared to young mice (Supplementary Fig. 1B, 3D). There was also prolonged Mmp8 elevation (Supplementary Fig. 1G). Aged mice exhibited no significant wound-induced increase in the expression of Tnf, Mcp1/Ccl2, Tgfb1, Mmp9 at the transcript level (Supplementary Fig. 1C, D, E, H). While Mmp3 mRNA increased in aged mice, this occurred later, peaking at day 12 (Supplementary Fig. 1F).

### Single-cell transcriptional atlas of day-6 murine wound beds from young and aged mice

To dissect the cellular heterogeneity and characterize specific cell populations contributing to age-dependent differences in wound healing, we performed single-cell RNA sequencing on dissociated wound tissue from young and aged mice harvested at day 6 post-wounding, a time point characterized by peak p16 induction and significant differences in wound healing kinetic. A total of 7,843 high-quality cells (4,145 from young, 3,698 from aged) were clustered using Seurat’s graph-based algorithm, revealing 20 transcriptionally distinct clusters (Fig. 3A an B, Supplementary Fig. 4). Based on canonical marker gene expression (Supplementary Fig. 4D-O), these clusters were annotated as diverse cell populations present in the wound microenvironment, including fibroblast, myeloid lineages (macrophages, monocytes, neutrophils), endothelial and lymphatic endothelial cells, T-cells, dendritic cells, erythroid cells, neural/glial cells, and adipocytes (Fig. 3B, Supplementary Fig. 4). UMAP visualization with cells colored by donor age confirmed cells from young and aged mice largely integrated within the same clusters (Fig. 3C). Analysis of cell distribution per cluster revealed general representation of both age groups across most clusters, although some clusters demonstrated modest shifts in proportional representation between young and aged wounds (Fig. 3D).

**Figure 3.**
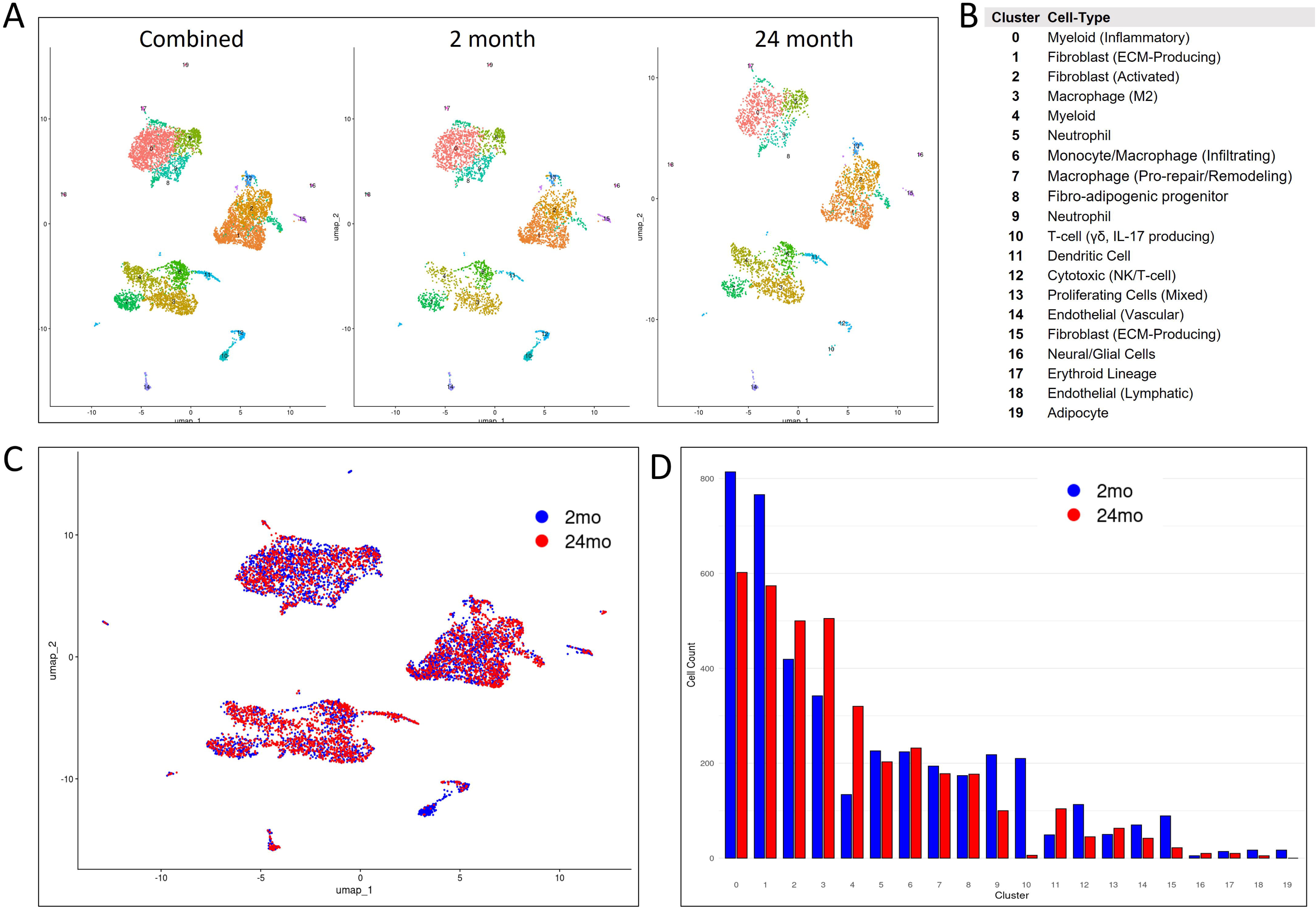
Single-cell transcriptional atlas of day-6 murine wound beds from young and aged mice. (A) UMAP projections of the integrated scRNA-seq dataset (total = 7,843 cells), including wound samples from 2-month-old (n = 4,145) and 24-month-old (n = 3,698) mice. Cells were normalized using SCTransform, integrated using Harmony to correct for age-related batch effects, and clustered with Seurat’s graph-based clustering algorithm (resolution = 0.3). Twenty transcriptionally distinct clusters (0–19) are visualized and color-coded by cluster identity. (B) Cell type annotations assigned to each cluster based on canonical marker gene expression. Identified populations include multiple fibroblast subsets, myeloid and neutrophil lineages, macrophages, endothelial and lymphatic cells, T-cell subsets, dendritic cells, erythroid cells, and adipocytes. (C) UMAP visualization with cells colored by age highlights effective batch correction and biological intermixing, with cells from 2-month-old mice shown in blue and from 24-month-old in red. (D) Bar plot demonstrating the distribution of cell numbers per cluster contributed by each age group. Blue bars represent cells from young mouse wounds; red bars represent cells from old mouse wounds. Y-axis denotes absolute cell counts (0–1,000 cells per cluster).

### Characterization of the defined senescent population in day-6 wound beds

We identified a specific senescent cell population based on combination of p16^+^ (Cdkn2a > 0), p21^+^ (Cdkn1a > 0), and Mki67^-^ (Mki67 = 0). Applying these criteria to the integrated day 6 dataset, we identified 234 cells meeting this definition, representing 2.98% of the total cells analyzed (Supplementary Fig. 5). UMAP visualization highlighted the distribution of these defined senescent cells across multiple clusters (Fig. 4A, B, Supplementary Fig. 5I). Analysis of the cellular composition of this defined senescent population revealed that the majority were fibroblasts (81 cells in Cluster 1, 87 in Cluster 2, 48 in Cluster 15), confirming fibroblasts as the predominant cell type exhibiting this senescent phenotype at day 6 (Fig. 4C). Specifically, Fibroblast Clusters 1 (ECM-Producing) and 2 (Activated) contained the highest numbers of senescent cells in both young and aged wounds, while Cluster 15 (ECM-Producing) was also a significant contributor (Fig. 6C). Other less abundant cell types within this defined senescent population included T-cells, neutrophils, macrophages, endothelial cells, dendritic cells, and adipocytes (Fig. 4C).

**Figure 4.**
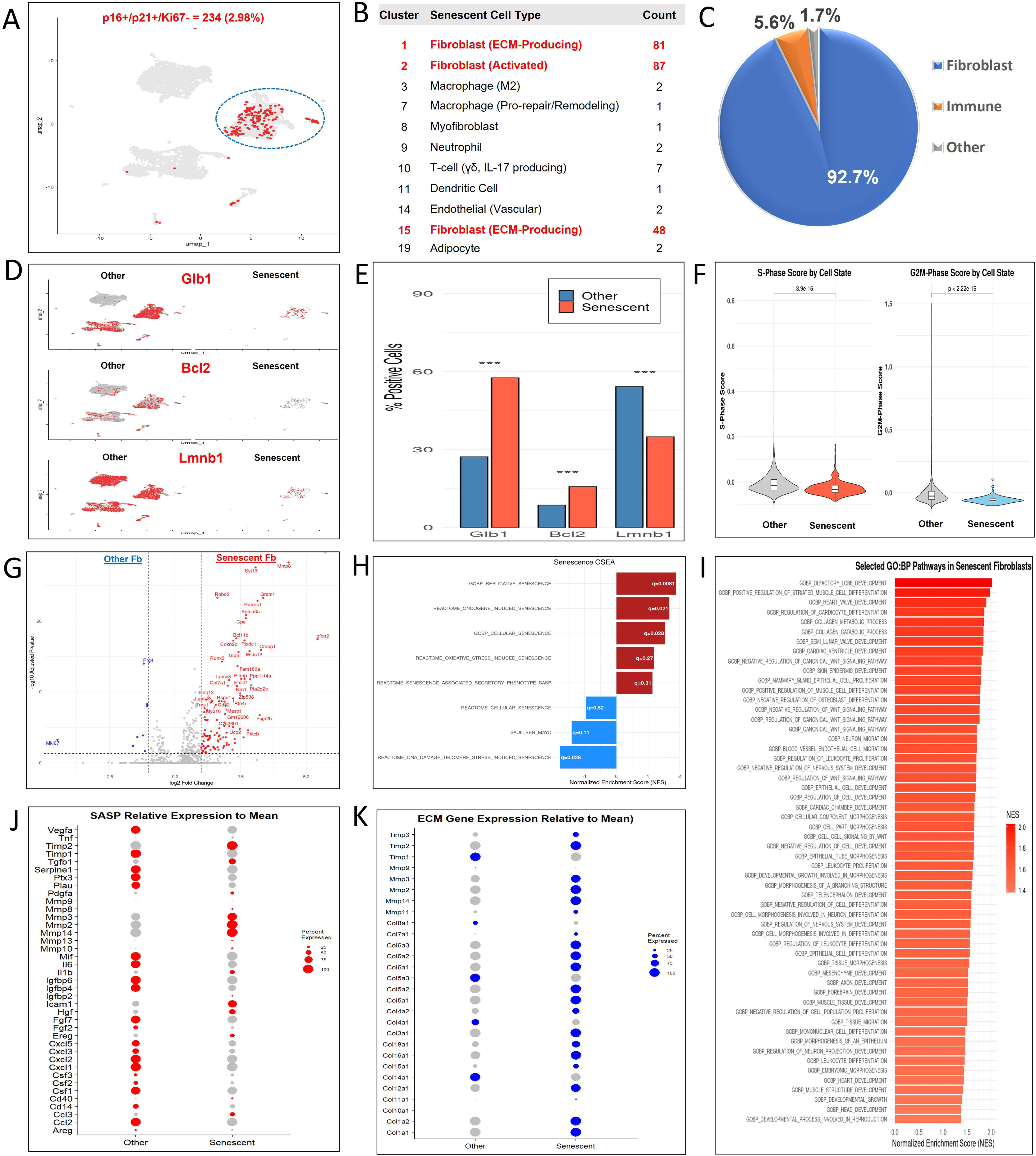
Multi-modal and functional validation characterizing p16+/p21+/Mki67-senescent wound cells in day-6 wound beds. (A) UMAP visualization of all integrated wound bed cells highlighting 234 p16+/p21+/Mki67-senescent cells in red (2.98% of total). (B) Cluster distribution of senescent cells across identified cell types. (C) Pie chart demonstrating the cellular composition of the senescent cell population. (D) Feature plots on the UMAP visualizing the expression of Glb1, Bcl2, and Lmnb1. (E) Bar plot comparing the percentage of positive cells expressing each indicated marker gene within the senescent and other cell populations. Statistical significance by Fisher’s exact test (*p<0.05, **p<0.01, ***p<0.001). (F) Violin plots displaying the S-phase and G2/M-phase cell cycle scores for senescent versus other cell populations, based on cell cycle gene expression. (G) Volcano plot of differentially expressed genes (logfc.threshold = 0.25, FDRq < 0.05) in senescent fibroblasts versus other fibroblasts. (H) GSEA of the senescence genesets with normalized enrichment score (NES) and FDRq-values (white). (I) Heatmap of selected significantly upregulated GO:BP pathways in senescent fibroblasts vs. other fibroblasts (FDRq<0.05). (J) Dot plot of SASP genes in senescent vs other fibroblasts. Dot size reflects percent expressing. (K) Dot plot of ECM genes in senescent fibroblasts. Dot size reflects percent expressing.

We further characterized the transcriptional profile of this defined senescent fibroblast population, delineated by the expression criteria p16^+^/p21^+^/Mki67^−^ in clusters 1, 2, 15. Quantitative comparison of the percentage of cells expressing specific marker genes revealed that the defined senescent population has significantly higher percentages of cells that demonstrate positive expression of Glb1 and Bcl2. Conversely, and consistent with senescent phenotype, the percentage of cells expressing Lmnb1 was significantly less frequent within this population (Fig. 4D and E). Evaluation of cell cycle status, inferred from S-phase and G2/M-phase scores, demonstrated significantly lower scores in the defined senescent population compared to other fibroblasts, consistent with a permanent state of cell cycle arrest (Fig. 4F).

The analysis of differential gene expression between senescent and non-senescent fibroblasts demonstrated a gain-of-function signature, with a predominant number of significant differentially expressed (DE) genes being up-regulated (such as Mmp8, Igfbp2, Grem1, and Sema3a). In contrast, only a few genes were found to be downregulated (see Fig. 4G), highlighting a transcriptionally active program in senescent fibroblasts. GSEA using senescence gene sets revealed positive enrichment for replicative, oncogene-induced, and general cellular senescence pathways, and negative enrichment for DNA-damage/telomere stress–induced senescence (NES and FDR < 0.05; Fig. 4H). GO BP analysis highlighted up-regulation of developmental, morphogenetic, and motility programs, including Wnt pathways, consistent with a dynamic pro-regenerative and ECM remodeling phenotype (Fig. 4I). At the SASP level, senescent fibroblasts overexpressed matrix regulators and growth factors (e.g. Igfbp2, Hgf, Cd40, Igfbp6, Icam1, Dcn, Tgfb1) but did not have higher expression of canonical inflammatory cytokines of SASP (Il6, Ccl3/4, Cxcl0, Il1b) compared to other fibroblasts (Fig. 4J). They also had higher expression levels of multiple collagen types alongside matrix metalloproteases than other fibroblasts, suggesting simultaneous ECM deposition and remodeling (Fig. 4K).

### Characterization of senescent fibroblasts and their pro-reparative phenotype is similar in human wounds

Analysis of publicly available single-cell RNA-seq data from acute human wounds (young donors in their 20s)^15^ (Fig. 5) identified the similar population of p16⁺/p21⁺/Mki67⁻ senescent fibroblasts transiently peaking at Day 7 and comprising 1.85 % of total wound cells (Fig. 5B, C). As in the mouse differential gene expression analysis, there was a predominance of upregulated genes compared to downregulated genes (Fig. 5E), reflecting a similar transcriptionally active, gain-of-function phenotype. Despite GSEA with established senescence signatures not indicating significant positive enrichment (data not shown), it did uncover notable negative enrichment for SAUL_SEN_MAYO (Fig. 5F). This pattern, akin to what is observed in mouse senescent cells, suggests a divergence from conventional senescence and points towards the presence of non-canonical senescent phenotype. Mirroring observations in young mice, young human senescent fibroblasts display a transcriptional profile marked by modest inflammatory output combined with a robust ECM remodeling signature. Specifically, senescent human wound fibroblasts had elevated levels of key collagen isoforms alongside matrix remodeling enzymes, and their inhibitors (Fig. 5G, H). GO:BP pathway enrichment further underscores this ECM-focused phenotype, demonstrating significant upregulation of collagen maturation and glycosylation pathways—hallmarks of intense matrix deposition and modification (Fig. 5I).

**Figure 5.**
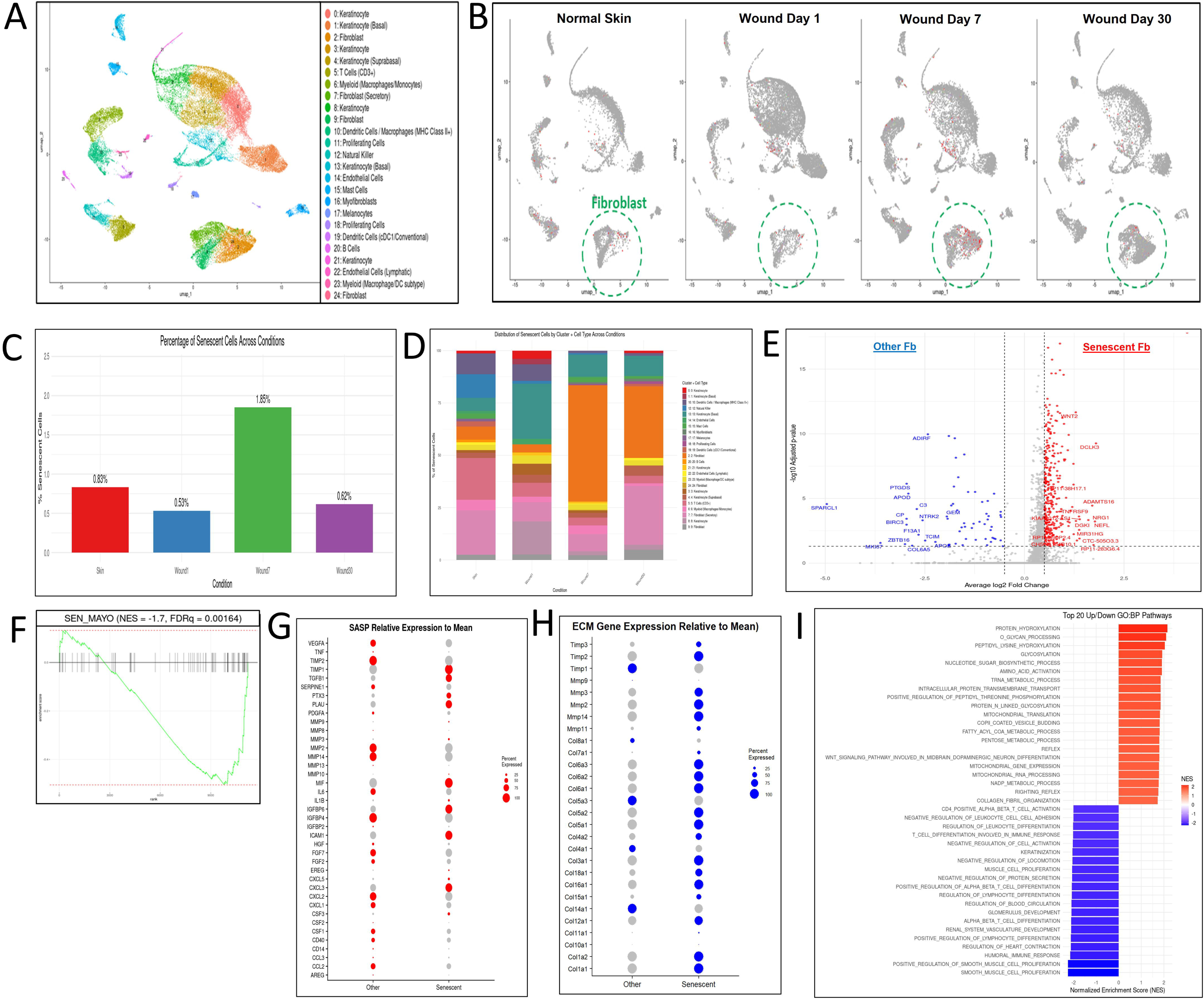
Transcriptional and functional characterization of senescent fibroblasts during human wound healing. (A) UMAP plot of all cell types from human skin and wound tissue (Skin, Wound day 1, Wound day 7, Wound day 30) annotated by cluster identity. (B) UMAP plots split by time point demonstrating the localization of senescent fibroblasts (p16⁺/p21⁺/MKI67⁻, red) across stages of wound healing. Fibroblast clusters are circled in green. (C) Bar graph quantifying the percentage of senescent cells among all cells across skin and wound stages. Senescent cells are most enriched at day 7 post-wounding. (D) Stacked bar plot demonstrating the cluster composition of senescent cells across each condition, indicating dynamic shifts in fibroblast senescent subtypes over time. (E) Volcano plot comparing gene expression in senescent vs. other fibroblasts at day 7, highlighting differentially expressed genes (red = upregulated in senescent cells). (F) GSEA plot demonstrating negative enrichment of the SenMayo senescence gene set in senescent vs. other fibroblasts at day 7 (NES = −1.7, FDR q = 0.0164). (G) DotPlot of SASP-related genes of senescent vs other fibroblasts at day 7. (H) DotPlot demonstrating expression of collagen-related genes in senescent vs. other fibroblasts at day 7. (I) Bar plot demonstrating the top 20 significantly enriched and downregulated GO:BP pathways in senescent fibroblasts at day 7, FDRq<0.05.

### Age-related differences in senescent wound fibroblasts

In aged mice, the proportion of senescent wound fibroblasts among all profiled cells was significantly lower (1.7%, 64 cells) compared to the young mice (4.1%, 170 cells) (Fig. 6A, B). Although the general distribution of senescent fibroblasts across clusters remained largely consistent between age groups, with clusters 1, 2, and 15 holding the highest percentages (Fig. 6C), aging did lead to subtle changes in cluster representation. Notably, there was a substantial decrease in Cluster 15, a key ECM producing fibroblast, in older mice.

**Figure 6.**
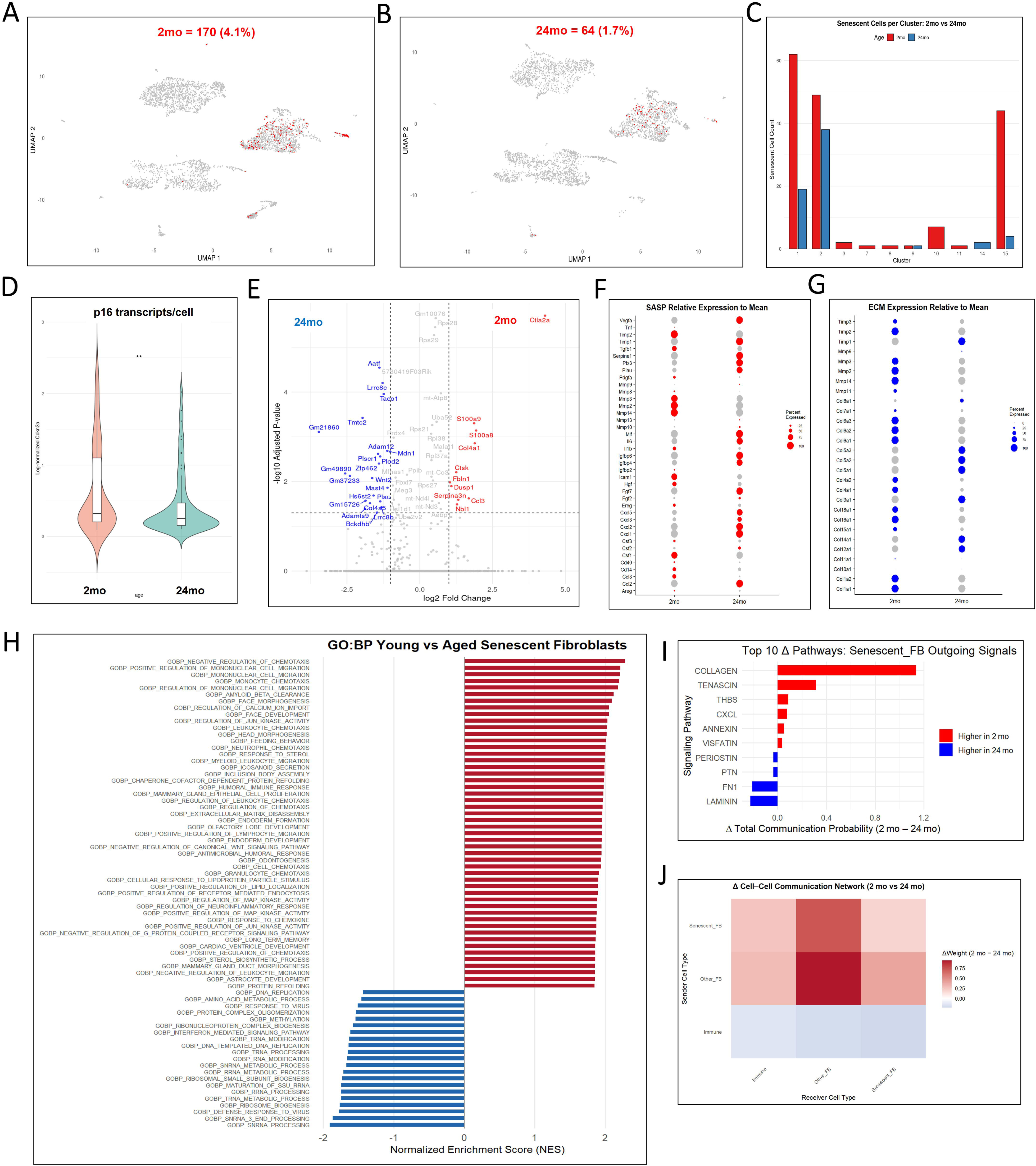
Age-dependent alterations in senescent wound fibroblast identity, secretome, and intercellular signaling. (A and B) UMAP projections of day 6 wound-bed cells, highlighting p16⁺/p21⁺/Mki67⁻ senescent fibroblasts (red) from 2-month-old (left; n = 170, 4.1%) and 24-month-old (right; n = 64, 1.7%) mouse wounds. (C) Cluster distribution of senescent fibroblasts. Bars represent per-cluster senescent cell counts in day 6 wounds from 2-month-old (orange) and 24-month-old (blue) mice. (D) Single-cell log-normalized Cdkn2a expression in day 6 wounds from 2-month-old versus 24-month-old mice, **p<0.01 Wilcoxon rank-sum test. (E) Volcano plot of differentially expressed genes in senescent fibroblasts (2 mo vs 24 mo); genes with |log₂ FC|>0.5 and adjusted p<0.05 are shown in red (up in 2 mo) or blue (up in 24 mo). (F) DotPlot of SASP-related genes in senescent fibroblasts from day 6 wounds in 2-month-old and 24-month-old mice. (G) DotPlot demonstrating expression of ECM-related genes in senescent fibroblasts from day 6 wounds in 2-month-old and 24-month-old mice. (H) GSEA of selected significant GO:BP pathways comparing senescent fibroblasts from day 6 wounds in 2-month-old and 24-month-old mice, FDRq<0.05. (I) CellChat plot demonstrating Δ total communication probability (2 mo – 24 mo) for Senescent_FB signaling pathways, with bar proportional to |Δ| and color indicating pathways stronger in young (blue) or aged (red) fibroblasts. (J). CellChat heatmap demonstrating Δ cell-cell communication (2mo – 24mo). Sender cell (y-axis): Receiver cells (x-axis). Red indicates enhanced cell-cell communication pairs in 2-month-old compared to 24-month-old mice.

In senescent wound fibroblasts from young mice, the median log₁ₚ₁-normalized Cdkn2a expression was 0.330 (IQR 0.208–1.07), compared to a median of 0.256 (IQR 0.155–0.494) in cells from old mice. On a linear scale (exp – 1), these correspond to roughly 0.39 versus 0.29 normalized transcript units, a ∼34 % increase in p16 levels in senescent fibroblasts from young mice (Fig. 6D). Notably, the broader expression range in cells from 2-month-old mice, including values exceeding 1.0, contrasts with the more constrained distribution in aged cells. This suggests that younger senescent fibroblasts not only have higher average Cdkn2a levels but also display greater heterogeneity and higher peak expression, indicating a more robust and varied senescence response.

Genome-wide differential expression analysis between young and aged senescent fibroblasts revealed mild number of significant differences on a volcano plot (Fig. 6E). However, targeted analysis of known SASP and ECM-related genes uncovered some age-dependent shifts. Young senescent fibroblasts exhibited elevated expression of genes involved in ECM remodeling (Mmps, Dcn, Timp2, Tgfb1), cellular growth and migration (Igfbp2, Hgf, Ereg), and inflammation/immune modulation (Cd40, Icam1). This gene expression profile is consistent with a more efficient and reparative wound healing phenotype in younger mice (Fig. 6H). In contrast, aged senescent fibroblasts demonstrated increased expression of pro-inflammatory factors, including Il6 and multiple Cxcl chemokines (Cxcl1, 2, 3, 5, 10), indicating a shift toward a more inflammatory secretome (Fig. 6F). Young senescent fibroblasts exhibited higher expression and prevalence of metalloproteinases (Mmp2, Mmp3, Mmp11, Mmp14) and fibrillar collagens (Col1a1, Col1a2, Col6a1, Col6a2, Col6a3). In contrast, aged senescent fibroblasts exhibited an increased proportion of cells expressing non-fibrillar and basement membrane collagens (Col4a2, Col5a1, Col5a2, Col12a1, Col14a1) (Fig. 6G).

GSEA comparing young and aged senescent fibroblasts revealed distinct age-associated functional signatures (Fig. 6H). Young senescent fibroblasts exhibited positive enrichment for developmental and morphogenetic pathways suggesting greater regenerative capacity or plasticity as well as pathways associated with stimulating cell-cell communication and activity with various cell types (Fig 6H). Conversely, aged senescent fibroblasts were more strongly enriched for interferon signaling and ribosomal and antiviral/interferon signaling, suggesting a dysfunctional, stressed state rather than a reparative one. (Fig. 6H).

CellChat analysis revealed shifts in outgoing signaling programs between young and aged senescent fibroblasts (Fig. 6I and J). In 2-month senescent fibroblasts, pathways involved in active ECM remodeling, particularly collagen, tenascin, and thrombospondin, and immune-cell communication – Cxcl chemokines demonstrated the largest positive Δ communication probabilities, indicating that young senescent fibroblasts more robustly broadcasting these matrix cues. In contrast, 24-month senescent fibroblasts displayed higher signaling through basement-membrane and adhesion molecules such as laminin and fibronectin (Fn1) (Fig. 6I). Correspondingly, analysis of the Δ cell–cell network (2 month – 24 month) confirmed that in 2 month wounds, senescent fibroblast → other fibroblast and senescent fibroblast → immune edges are strongly elevated, forming a dense fibroblast-centric communication hub; by 24 months, that hub collapses and immune cells assume a relatively stronger sender role, indicative of chronic inflammation rather than repair (Fig. 6J).

In sum, young senescent fibroblasts mount a pro-repair chemokine and ECM-remodeling program and broadcast it to their neighbors, whereas aged senescent fibroblasts lose that reparative signaling and instead exhibit proteotoxic/inflammatory features that fail to promote efficient healing.

## Discussion

The transient induction of cellular senescence during acute wound healing is increasingly recognized as an important biological process facilitating normal tissue repair^8,9^. Building upon prior studies demonstrating transient upregulation in p16, p21, SA-β-gal, and certain SASP factors during the physiological wound healing process^8,9,16^, we employed a comprehensive and orthogonal multi-modal approach, integrating histology, qPCR, Western blot, ELISA, and single-cell RNA sequencing, to investigate the dynamics of senescence induction and the cellular and molecular characteristics of senescent populations during full-thickness cutaneous wound healing in young and aged mice. Our findings demonstrate that the wound-induced senescence response is robust and kinetically coordinated in young mice but significantly attenuated and dysregulated in aged animals. This impairment in acute senescence induction in aged wounds correlates with the observed delay in macroscopic wound closure kinetics, providing evidence that a compromised transient senescence response may mechanistically contribute to age-related healing deficits.

Distinct age-dependent patterns of senescence marker and SASP factor expression were revealed in our bulk wound tissue analyses. While young mice exhibited coordinated transient peaks of p16, p21, p53, SA-β-gal, and multiple SASP factors (Il6, Mcp1/Ccl2, Mmp3, Mmp9, Tnf, Tgfb), aged mice demonstrated a striking failure to significantly induce many of these markers above their baseline levels. The bulk tissue SASP profile in aged wounds was not only deficient for several key factors (Mcp-1/Ccl2, Mmp9, Tgfb) but also displayed prolonged expression of certain pro-inflammatory mediators such as Il6 and Mmp8, contributing to a prolonged inflammatory state. This aligns with the concept of “inflammaging,” where dysregulation of inflammation is a hallmark of aging, and suggests that the age-related delay in healing is associated with both a failure to induce a beneficial transient response and a shift towards a more detrimental inflammatory state that impedes resolution and repair^17^. Importantly, because senescent cells are relatively rare in whole-tissue samples, not all detected SASP factors can be attributed solely to those cells.

Our single-cell RNA sequencing analysis provided critical resolution into the cellular sources and characteristics of the senescence response at day 6, a pivotal time point when transition from inflammatory to proliferative phase occurs^18^. We successfully identified and characterized a specific p16^+^/p21^+^/Mki67^−^ senescent cell population based on canonical marker expression at the single-cell level, confirming that fibroblasts constitute the most abundant cell type exhibiting this phenotype in murine wounds at this stage. These p16⁺/p21⁺/Mki67⁻ senescent fibroblasts represented a much smaller fraction of all wound cells in aged mice than in the young, and their per-cell p16 expression was lower, indicating both fewer senescent cells and a weaker p16 signal in the aged wounds. This diminished presence of senescent cells in aged wounds likely represents a fundamental defect in the number of cells entering this state or maintaining this beneficial phenotype with aging.

Beyond their reduced numbers, aged wounds harbor senescent fibroblasts with a fundamentally altered transcriptional program. In 2-month-old mice, senescent fibroblasts express high levels of reparative and ECM-remodeling factors—Igfbp2, Hgf, Cd40, Mmp14, Icam1, Mmp3, Mmp2, Dcn, Timp2, Ereg, Tgfb1—along with abundant MMPs (Mmp2, Mmp3, Mmp11, Mmp14) and fibrillar collagens (Col1a1, Col1a2, Col6a1, Col6a2, Col6a3). GSEA further confirms enrichment for morphogenetic and matrix-building programs with increased cell communication activity (Fig 6H). By contrast, senescent fibroblasts from 24-month-old mice up-regulate pro-inflammatory mediators (Il6, multiple Cxcl chemokines) and more frequently express non-fibrillar/basement-membrane collagens (Col4a2, Col5a1, Col5a2, Col12a1, Col14a1). Their signature shifts toward interferon signaling and proteostastic stress (Fig 6H) which has been reported in aged, dysfunction dermal fibroblasts^19,20^. Complementing these gene-level shifts, CellChat demonstrates that young senescent fibroblasts predominantly broadcast through collagen and tenascin pathways, consistent with active ECM rebuilding, whereas aged cells switch to fibronectin and laminin signaling, favoring matrix stabilization, stronger adhesion, and inflammatory cross-talk. In sum, senescent fibroblasts in young wounds drive a pro-regenerative, ECM-focused SASP, while aged senescent fibroblasts adopt a pro-inflammatory, matrix-stabilizing phenotype that likely impedes proper wound resolution. Furthermore, young senescent fibroblasts mount a pro-repair chemokine and ECM-remodeling program and broadcast it to their neighbors, whereas aged senescent fibroblasts lose that reparative signaling and instead exhibit proteotoxic/inflammatory features that fail to promote efficient healing.

Our analysis of publicly available human wound scRNA-seq data^15^ provides strong translational support for our murine findings. Using the same p16⁺/p21⁺/MKI67⁻ definition, we identified a transient senescent fibroblast population peaking at Day 7 of human wound healing. These cells exhibited a pro-reparative transcriptional profile enriched for ECM organization, closely mirroring the phenotype of young murine senescent wound cells. This conservation across species suggests a conserved core senescence program critical for effective acute wound healing. Although the human dataset lacked age-matched samples for direct comparison, independent studies support age-related deficits in wound-induced senescence. One study comparing young (<30 years) and older (>75 years) individuals revealed upregulation of p21, p53, and MMP-9 after injury only in young subjects, with a non-significant trend for p16 (p=0.08) and no induction in aged skin (p=0.25)^9^. While limited by sample size and lack of cell-type resolution, these findings align with our observation that aging disrupts the normal transient wound-induced senescence response and may contribute to impaired wound repair in humans.

We hypothesize that age-related delays in wound healing are related to two distinct senescent cell populations with (1) increases in resident chronic senescent cells present in aged skin^21,22^, and (2) decreases in acutely induced senescent cells that emerge in response to injury. Chronic senescent cells may impair repair by creating a pro-inflammatory, non-regenerative environment^23,24^. We observed elevated baseline expression of p16, p21, and SA-β-gal activity in unwounded aged skin, consistent with prior studies in both mice and humans. Importantly, our previous work demonstrated that experimentally increasing senescence in young skin via transplantation of irradiation-induced senescent fibroblasts delayed healing^5^, while clearing senescent cells with topical senolytic in aged mice improved wound closure^25^. These findings suggest that chronic senescent cells hinder, while acute senescent cells support, wound healing, and that imbalances between these populations in aging may underlie impaired repair. The second population of senescent cells responsible for transient upregulation of senescence signaling during wound healing is robust only in young individuals, and lack of this transient upregulation may be implicated in delayed wound healing in aging given pro-reparative, ECM-producing and remodeling phenotype of these cells.

The precise reasons for the absence of this crucial transient wound-induced upregulation of acute senescence signaling during aged wound healing remain unknown. Understanding the unique roles and characteristics of these distinct cell populations, particularly the mechanisms governing their induction, maintenance, and functional phenotype with age, is crucial for devising targeted strategies to enhance tissue repair in aging individuals. Future investigations aimed at dissecting the precise molecular triggers of age-related senescence dysregulation in wounds and exploring therapeutic strategies aiming at restoration of the physiological transient senescence response or modulating the phenotype of senescent cells in aged wounds hold significant promise for improving healing outcomes in the older adult population.

While our study adds to the framework of age-related differences in wound-associated senescence at single-cell resolution, we acknowledge several limitations. Our analysis focused on a single time point (Day 6), which coincides with peak p16 expression but does not capture the full temporal dynamics of senescence throughout wound healing; longitudinal single-cell studies will be important to define the initiation, resolution, and functional transitions of senescent cells over time. Although we used p16⁺/p21⁺/Mki67⁻ signature as a marker of senescence, it may not encompass the full heterogeneity of senescent cell states as other subtypes or marker combinations could also contribute to repair or pathology. Our conclusions are drawn from discrete comparisons between two age groups in scRNA-seq data, which may miss subtle, continuous shifts in gene expression or rarer senescent phenotypes. Finally, while single-cell transcriptomes reveal cell-intrinsic programs, they do not directly measure protein activity, secreted factors, or cell–cell interactions *in situ*. Despite these caveats, our findings establish a foundation for understanding how aging reshapes the transient senescence response and highlight the therapeutic potential of targeting specific senescent cell phenotypes to improve wound healing in older adults.

## Conclusion

Aging significantly delays cutaneous wound healing, coinciding with an impaired and dysregulated transient senescence response. In young wounds, p16⁺/p21⁺/Mki67⁻ senescent fibroblasts exhibit a robust, pro-reparative phenotype marked by ECM remodeling and matrix-modulating SASP factors. In contrast, aged wounds demonstrate a reduced percentage of these cells and a shift toward a pro-inflammatory, less regenerative profile. These findings highlight an important role of transient, fibroblast-driven senescence in promoting effective wound repair, one that becomes blunted and maladaptive with age. Targeting this dysfunction offers a promising therapeutic avenue to enhance healing in older adults but also underscores the need for caution, as senolytic therapies move forward, to avoid disrupting beneficial acute senescence responses essential for tissue regeneration.

## Data availability statement

The raw and processed single-cell RNA sequencing data generated for this study have been deposited in the NCBI Gene Expression Omnibus (GEO) repository and are publicly available under accession number [GEO accession number will be inserted here upon deposition prior to publication]. All other data supporting the findings of this study are available within the article and its supplementary materials or from the corresponding author upon reasonable request.

## Author contribution

DSR designed and supervised the study and completed computational analysis. MS wrote the original draft of the manuscript, performed and analyzed qPCR experiments, performed and analyzed Western blot and ELISA experiments, performed FACS sorting, and performed statistical analysis. RJRST analyzed wound imaging data. MS and RJRST performed and analyzed IVIS imaging experiments. MS, RJRST, JD, QW, and GHS completed wound healing experiments, performed and analyzed immunostaining and SA-β-gal staining. All authors reviewed, revised, and approved the final version of the manuscript.

## Supplemental figure legends

**Supplementary Figure 1. Additional senescence and SASP markers during wound healing in young and aged mice.** qRT-PCR analysis of wound tissue from young and aged mice at different time points during wound healing. Expression of (A) p53, (B) Il6, (C) Tnf-α, (D) Mcp-1, (E) Tgf-β, (F) Mmp3, (G) Mmp8, and (H) MMP-9 is shown relative to β-actin expression. N=5 per age group per time point. Asterisks (*) indicate statistically significant differences (p<0.05).

**Supplementary Figure 2. p21-positive cells transiently accumulate during young but not during old wound healing.** (A and B) Representative immunofluorescence images demonstrating p21 (red) and DAPI (blue) staining in wound tissue from young (A) and aged (B) mice at 0, 6, 12, 18, and 24 days post-wounding. The epidermis (epi) and dermis (der) are indicated for reference. (C) Quantification of the percentage of p21-positive cells in wound tissue from young and old mice over the wound healing time course. (D) qRT-PCR analysis of p21 gene expression in wound tissue from young and aged mice over the wound healing time course, shown as relative expression to β-actin. N=5 per age group per time point. Asterisks (*) indicate p<0.05.

**Supplementary Figure 3. Protein expression confirmation of senescence and SASP markers from young and old mouse wounds.** Analysis of wound tissue protein lysates from young and aged mice during wound healing. (A) Representative Western blots demonstrating protein levels of MMP-9 and p21 at 0, 6, 12, 18, and 24 days post-wounding. GAPDH is included as a loading control. (B) Quantification of relative MMP-9 protein levels over time in young and old mice. Protein levels relative to GAPDH from Western blots shown in (A). (C) Quantification of relative p21 protein levels over time in young and old mice. Protein levels relative to GAPDH from Western blots shown in (A). (D) Quantification of IL-6 protein levels in wound tissue lysates from young and aged mice over the wound healing time course, measured by ELISA. N=5 per age group per time point. Asterisks (*) indicate p<0.05, and double asterisks (**) indicate p<0.01.

**Supplementary Figure 4: Single-cell RNA sequencing analysis of day 6 wound tissue.** (A-C) UMAP plots of single-cell RNA sequencing data from Day 6 wound tissue. Cells are colored by unsupervised clustering. (A) Combined data from 2-month and 24-month-old mice. (B) Data from 2-month-old mice only. (C) Data from 24-month-old mice only. Cluster numbers are indicated, and the legend in (A) and (C) demonstrates the corresponding cluster identities based on marker gene expression. (D-O) Feature plots demonstrating the expression levels of selected marker genes overlaid on the combined UMAP from (A). Gene names are indicated above each plot. Color intensity represents the normalized expression level of the gene, with darker shades indicating higher expression. These markers were used to assist in the identification and annotation of cell clusters. (D) Col1a1, (E) Adipoq, (F) Ptprc, (G) Cd68, (H) S100a6, (I) Pecam1, (J) Cd3d, (K) Hba-a1, (L) Itgam, (M) Pdgfra, (N) Mki67, (O) Ly6c1.

**Supplementary Figure 5: Expression analysis of senescence and proliferation markers in wound tissue.** (A-I) UMAP plots illustrating the distribution of cells expressing specific senescence and proliferation markers. Cells are colored based on whether they are negative (grey) or positive (red) for the indicated gene or combination of genes. The plots demonstrate cells positive for: (A) Cdkn2a, (B) Cdkn1a, (C) Co-expression of Cdkn2a and Cdkn1a (Cdkn2a > 0 & Cdkn1a > 0), (D) Cdkn2a expression in proliferating cells (Cdkn2a > 0 & Mki67 > 0), (E) Cdkn1a expression in proliferating cells (Cdkn1a > 0 & Mki67 > 0), (F) Co-expression of Cdkn2a and Cdkn1a in proliferating cells (Cdkn2a > 0 & Cdkn1a > 0 & Mki67 > 0), (G) Cdkn2a expression in non-proliferating cells (Cdkn2a > 0 & Mki67 = 0), (H) Cdkn1a expression in non-proliferating cells (Cdkn1a > 0 & Mki67 = 0), (I) Co-expression of Cdkn2a and Cdkn1a in non-proliferating cells (Cdkn2a > 0 & Cdkn1a > 0 & Mki67 = 0). (J) Table summarizing the number and percentage of cells falling into the defined expression categories shown in the UMAP plots. The ‘Condition’ column indicates the expression criteria, ‘Count’ demonstrates the count of cells meeting those criteria, and ‘Percent’ demonstrates the percentage of the total cells analyzed represented by that group.

## Notes

**Funding information:** This work was supported by grants from the National Institute of Aging [R03AG067983, K76AG083300]; Laszlo N. Tauber Professorship in Surgery.

**Disclosure:** the authors declare no conflicts of interest.

### Competing Interest Statement

The authors have declared no competing interest.

