## Supplementary table 1 for "Diminished and altered cellular senescence response in delayed wound healing of aging"

### Supplementary data

**Table S1. Primer sequences used.**

| Gene | Forward primer sequence (5'-3') | Reverse primer sequence (5'-3') |
|---|---|---|
| p16 | TGTTGAGGCTAGAGAGGATCTTG | CGAATCTGCACCGTAGTTGAGC |
| p21 | CTGAGCGGCCTGAAGATTCC | CCAATCTGCGCTTGGAGTGA |
| p53 | ACGCTTCTCCGAAGACTGG | TCCATGCAGTGAGGTGATG |
| Il-6 | TACCACTTCACAAGTCGGAGGC | CTGCAAGTGCATCATCGTTGTTC |
| MCP-1 | GCTACAAGAGGATCACCAGCAG | GTCTGGACCCATTCCTTCTTGG |
| MMP-3 | CTCTGGAACCTGAGACATCACC | AGGAGTCCTGAGAGATTTGCGC |
| MMP-8 | GATGCTACTACCACACTCCGTG | TAAGCAGCCTGAAGACCGTTGG |
| MMP-9 | ACGACATAGACGGCATCCA | GCTGTGGTTCAGTTGTGGTG |
| TNF-α | GATCGGTCCCCAAAGGGATG | CCACTTGGTGGTTTGTGAGTG |
| TGF-β | ACTGGAGTTGTACGGCAGTG | GGGGCTGATCCCGTTGATTT |
| β-actin | GCACTGTGTTGGCATAGAGG | GTTCCGATGCCCTGAGGCTCTT |
