## Supplementary figures and images for "Diminished and altered cellular senescence response in delayed wound healing of aging"

### Supplementary figure 1

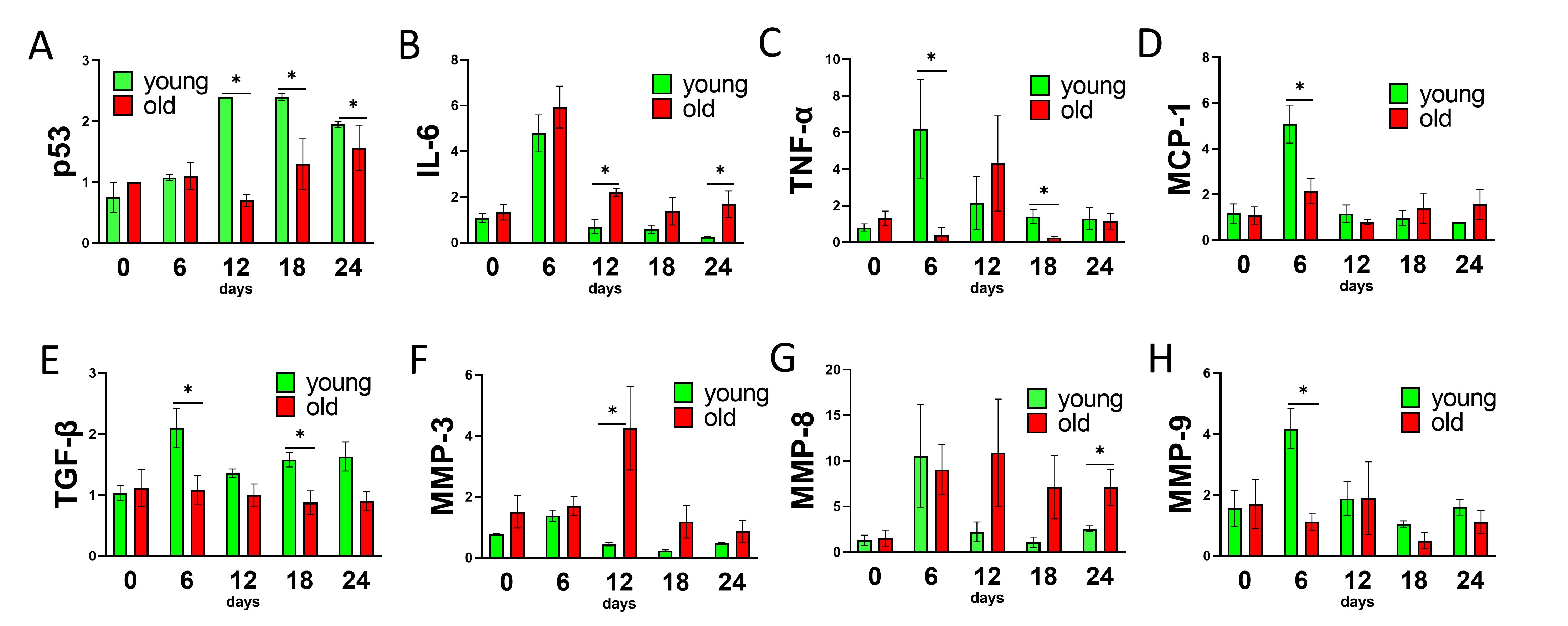

### Supplementary figure 2

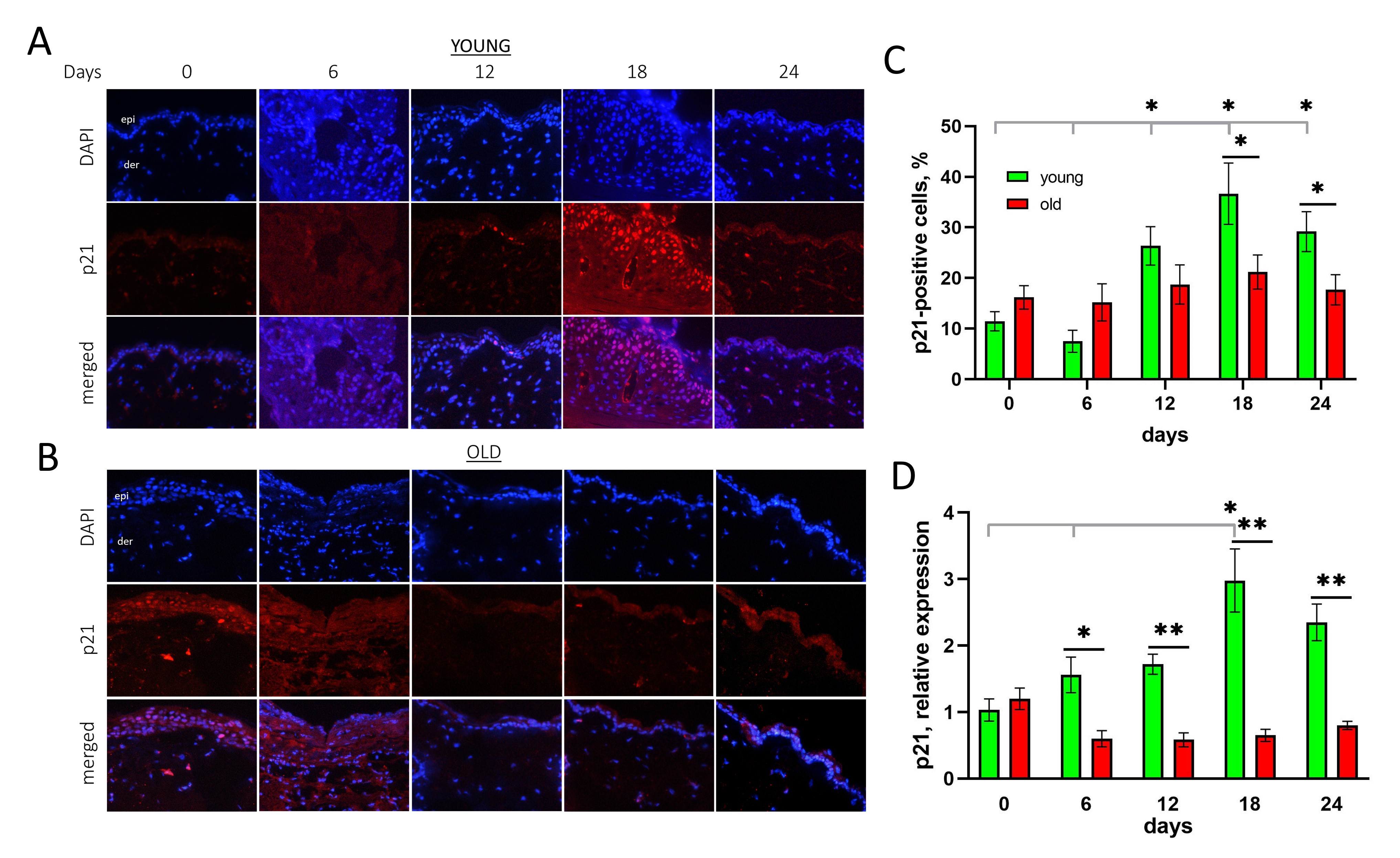

### Supplementary figure 3

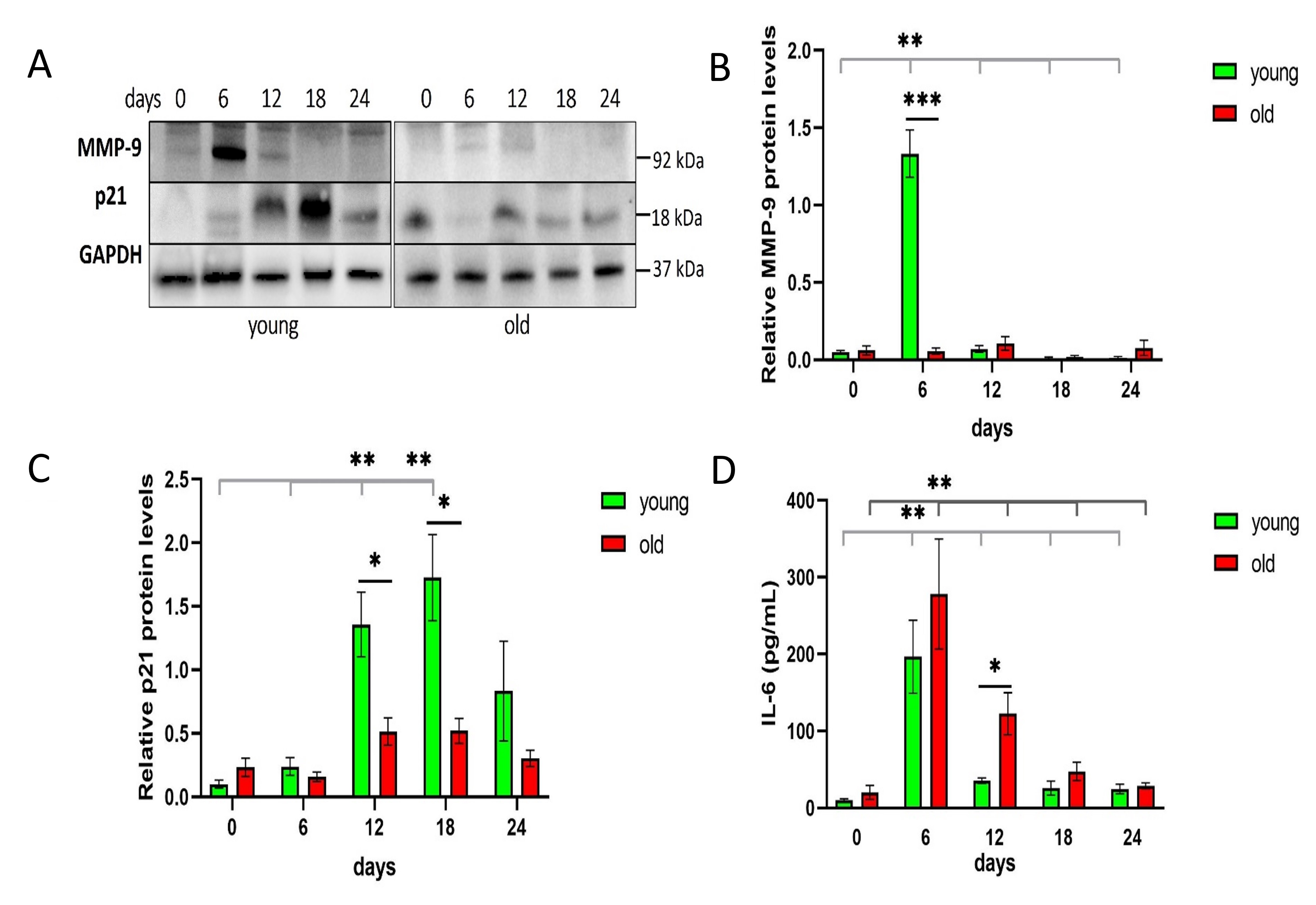

### Supplementary figure 4

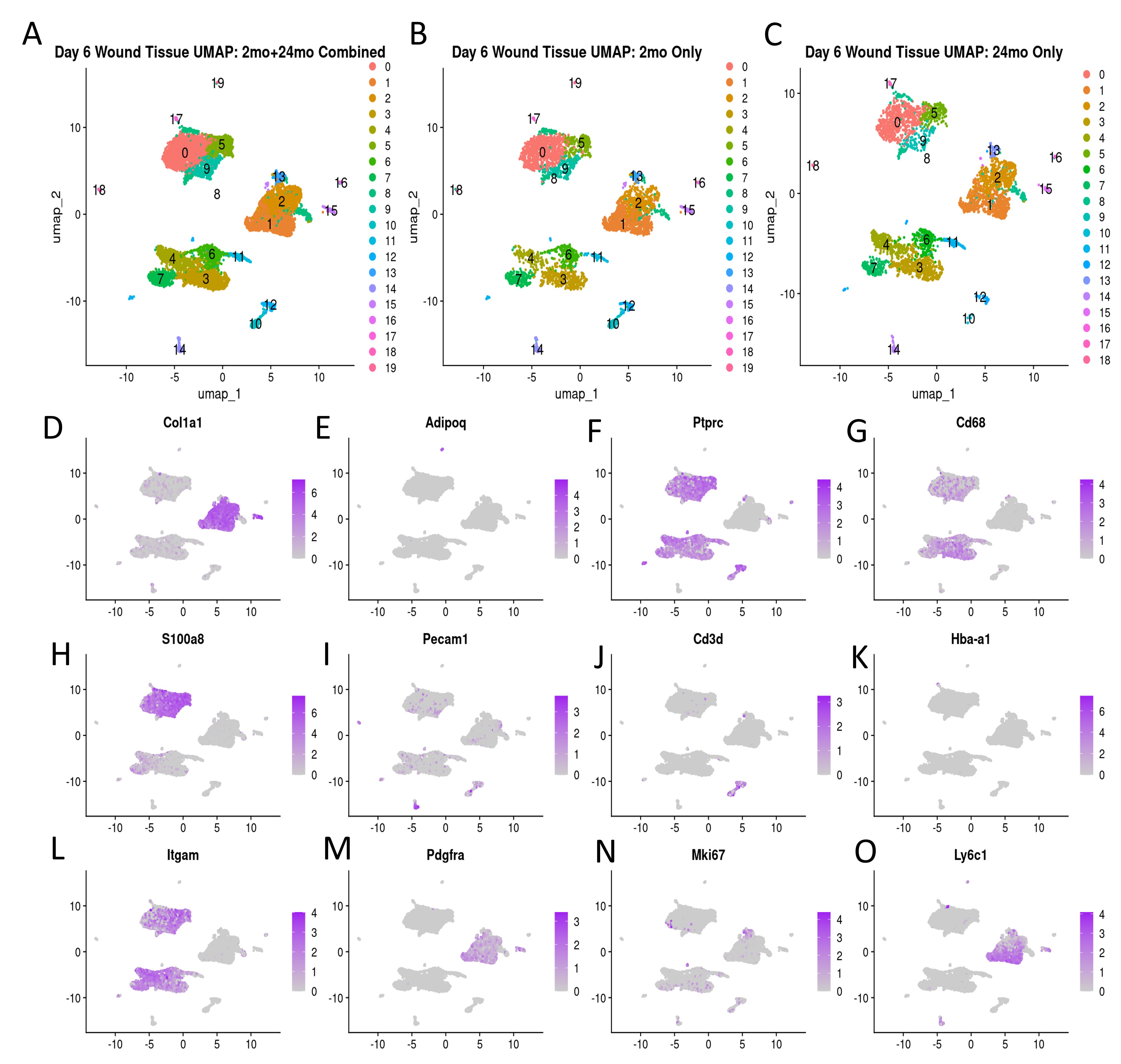

### Supplementary figure 5

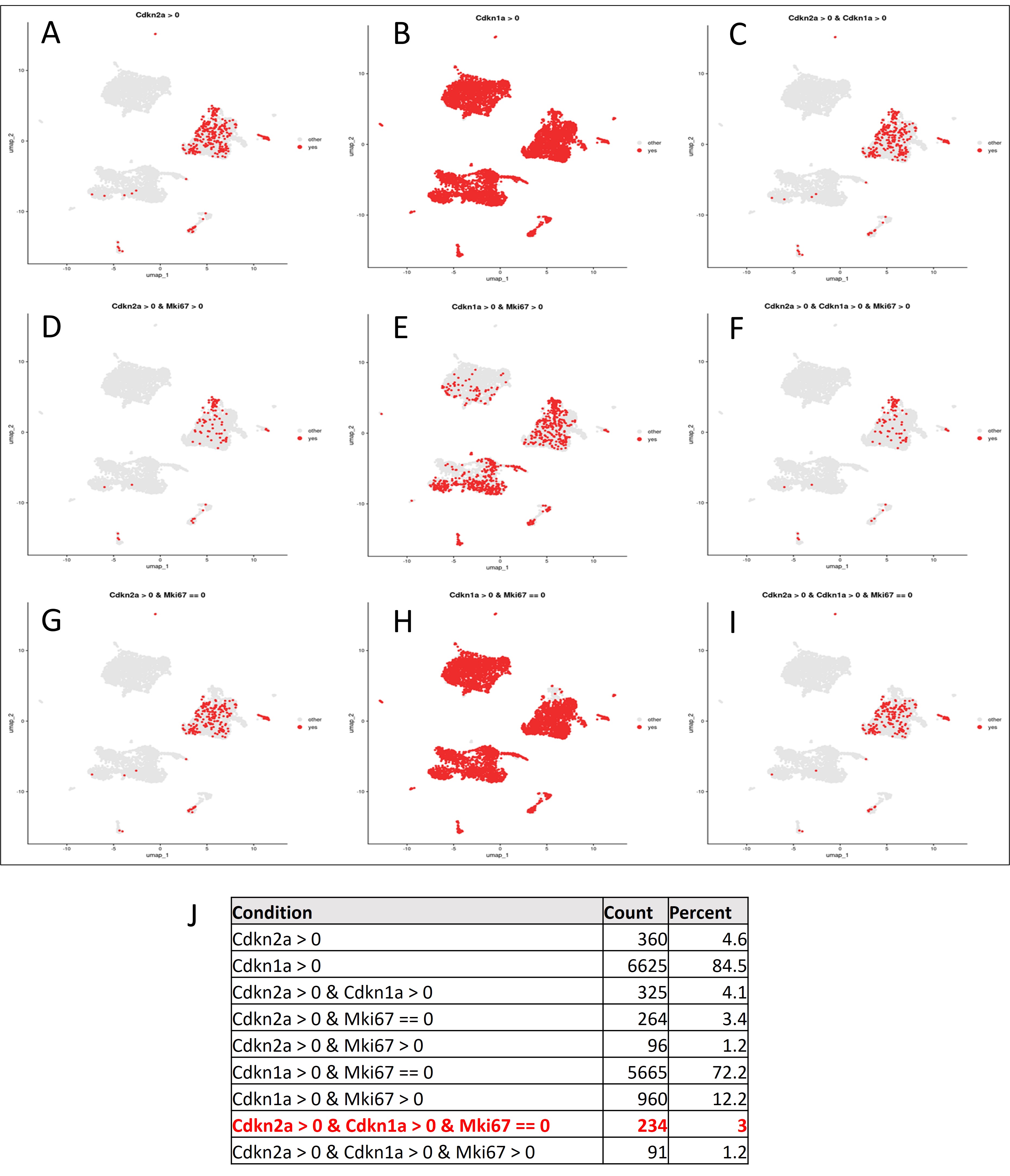
